# Reversing Nicotine Toxicity: how Platelet-Rich Plasma Enhances Cell Recovery through Autophagy Modulation

**DOI:** 10.1101/2025.10.06.680686

**Authors:** Julie Vérièpe-Salerno, José Antonio Cancela, Solange Vischer, Antoine Turzi, Muriel Cuendet, Catherine Giannopoulou, Sarah Berndt

## Abstract

**Background:** Chronic exposure to nicotine significantly exacerbates periodontitis, a prevalent inflammatory disease, by inducing cellular processes such as autophagy and inflammation in gingival fibroblasts. Current therapies often fail to fully address these cellular alterations in smokers, highlighting a need for innovative therapeutic and generative approaches.

**Objective:** This study explores the therapeutic potential of Platelet-Rich Plasma (PRP), a blood-derived product, to modulate nicotine-induced biological activities in primary gingival fibroblasts. It aims at understanding the underlying cellular mechanisms and assessing the efficacy of PRP as an adjunct treatment for periodontitis in smokers.

**Methods:** Gingival fibroblasts were treated with increasing concentrations of nicotine, which led to senescence and autophagy. Subsequent treatment with autologous PRP was evaluated for its effect on the reversion of these processes, by measuring cell migration and proliferation, metabolic activity, as well as by looking at senescence and autophagic markers. A *Caenorhabditis elegans* model of autophagy was used to assess nicotine and PRP biological activities in an *in vivo* environment.

**Results:** Nicotine at high concentrations triggered cellular vacuolization, a decrease in metabolism, viability and proliferation that was partially (with 500 ng/ml nicotine) or completely (with 250 ng/ml nicotine) reversed by a concomitant treatment with 10% PRP. Nicotine alone (250 ng/ml) slightly enhanced migration, while concomitant treatment between nicotine and PRP significantly increased their migration potential. In *Caenorhabditis elegans*, PRP reduced the nicotine-induced autophagic activity, as evidenced by decreased numbers of autophagosome and a higher number of viable worms during adulthood in comparison to nicotine control conditions. A screening of gingival fibroblast secretome revealed a modulation of autophagy-related cytokines in response to nicotine and/or PRP.

**Conclusion:** The findings demonstrate that PRP could effectively inhibit nicotine-induced autophagy gingival fibroblasts, offering insights into its possible use as a therapeutic tool for managing periodontitis in smokers. The study underscores the potential of PRP in altering disease progression by modulating key cellular processes affected by smoking.

## Introduction

Periodontitis is an inflammatory disease characterized by microbially associated host-mediated inflammation that results in loss of periodontal attachment. Bacterial biofilm formation initiates gingival inflammation. Periodontitis occurs when the balance between the microbial biofilm and the host is lost, due to dysbiosis that induces a perturbation of the host homeostasis in susceptible individuals. Current evidence supports that periodontitis is a multifactorial disease, as several systemic and environmental factors may modify the host’s susceptibility to periodontitis and the disease’s extent, severity, progression, and response to therapy [1].

Tobacco smoking has been identified as a major risk factor for periodontitis. The damaging effect of smoking has been shown in several epidemiological and clinical studies. Furthermore, studies on the potential mechanisms by which smoking tobacco may predispose to periodontal disease have been conducted. It appeared that smoking may affect the vasculature, the humoral immune system, and the cellular immune and inflammatory systems [2]. For example, neutrophil migration and chemotaxis in the periodontal tissues were impaired in smokers, the vasculature was decreased, and several functions of gingival and periodontal ligament fibroblast were impaired [3-5].

Nicotine is highly harmful to oral cells and significantly contributes to the progression of periodontal disease. It exhibits cytotoxic effects on oral epithelial cells and fibroblasts, which are critical for maintaining the structure and function of gum tissues. Nicotine reduces cell viability and impairs the proliferation of these cells, limiting the body’s ability to repair damaged tissues. Moreover, it intensifies the inflammatory response within the oral cavity, leading to the release of pro-inflammatory cytokines that cause further tissue and bone damage. Another key issue is that nicotine induces vasoconstriction, which restricts blood flow to the gums, thereby reducing oxygen and nutrient delivery. This impaired circulation slows down healing processes and increases susceptibility to infections [6].

In the context of periodontitis, nicotine accelerates alveolar bone loss, which compromises tooth support. This is primarily driven by nicotine’s effect on osteoclast activity, promoting the breakdown of bone tissue. Additionally, nicotine hampers the immune system’s ability to manage bacterial plaque, which allows periodontal pathogens to proliferate unchecked, thereby worsening the disease. It also alters the oral microbial environment, favoring the growth of harmful bacteria associated with periodontitis. Furthermore, nicotine interferes with collagen production, which is essential for gum and bone integrity. It reduces collagen synthesis and increases its degradation, leading to weaker gums and the formation of deeper periodontal pockets [7].

Nicotine also significantly impacts wound healing and tissue regeneration in periodontal procedures. The substance reduces fibroblast activity, impairs blood supply, and generally delays the healing of periodontal tissues following interventions like scaling and root planning. These combined effects highlight why individuals who smoke or use nicotine products are at a considerably higher risk for developing severe periodontitis and face greater challenges in managing the disease.

Autophagy is a cellular process that allows cells to degrade and recycle their own components, helping to maintain cell homeostasis and survival during stress conditions. The term “autophagy” comes from Greek, meaning “self-eating.” It involves the formation of a double-membrane structure called the autophagosome, which engulfs damaged organelles, proteins, and other cellular debris. The autophagosome then fuses with a lysosome, forming an autolysosome, where the acidic environment and lysosomal enzymes work together to degrade the contents, which are then either recycled for energy or repurposed to build new cellular components. Autophagy also plays a role in immune responses, development, and adaptation to nutrient deprivation. Dysregulation of autophagy is linked to various diseases, including cancer, neurodegenerative disorders like Alzheimer’s disease or amyotrophic lateral sclerosis, and metabolic diseases such as diabetes [8].

Periodontal treatment aims to reduce the inflammatory response, decrease disease progression, and ideally, regenerate lost periodontal structures. Thus, in addition to meticulous mechanical cleaning of the tooth surfaces, several regenerative techniques have been introduced for the treatment of periodontitis to remove the bacterial deposits (plaque and calculus). Among these, platelet-rich plasma (PRP) has been proposed due to its antibacterial properties, and its role in homeostasis and wound healing [9]. Several *in vitro* studies have shown a significant effect of PRP on several functions including osteoblast and fibroblast proliferation and differentiation [10-13]. Recent clinical research confirmed that using PRP as an agent for a periodontal wound increased the concentration of growth factor locally, thus enhancing the healing outcome [14]. Platelet concentrates (PCs) are easily obtained from the patient blood by centrifugation and are widely used for sinus floor elevation, alveolar ridge preservation, periodontal bone defects, guided bone regeneration, and treatment of gingival recession. Autologous PCs are ideal for this purpose because they have a high concentration of platelets, growth factors, and cytokines [15].

The aim of the present research project was to investigate the influence of PRP on various cellular functions of nicotine-treated gingival fibroblasts (GF). We showed that the PRP can partially compensate for the negative effect of nicotine on the cells of the periodontium, thus it could be recommended in the treatment of periodontitis in smokers in a preventive or curative approach.

## Materials and methods

### Cell cultures

Human GF were obtained from biopsies of donors clinically treated for various reasons. After washing with Dulbecco’s Phosphate Buffered Saline solution (DPBS, Gibco) to dilute the oral bacterial flora of the gingival tissue, small fragments of biopsy (2 mm square) were plated in culture dishes in control growth medium (GM): DMEM (Capricorn Scientific) containing 10% heat-inactivated fetal bovine serum (FBS, Capricorn Scientific) and 1% penicillin-streptomycin (Pan Biotech). Dishes were incubated at 37°C in 5% CO_2_ humidified atmosphere for 48 h, undisturbed. After 48 h, the medium was replaced. Two different donors were used to conduct all the experiments. The procedure conformed to the principles of the Declaration of Helsinki and was approved by CCER Geneva ethics committee (ID 2017–00700, approved on 18th October 2018). GF were used between passages 2 to 10.

### Preparation of human allogenic PRP and PPP

Allogenic blood samples from donors were processed into specific medical devices (RegenKit-BCT, Regen Lab SA). These devices produce a standardized leucocyte poor PRP thanks to a separator gel that isolates the plasma and the platelets from the other blood components. Ten ml of blood were collected per tube and centrifuged at room temperature for 5 min in a Regen Lab centrifuge (Drucker Horizon-6-FA) at a force of 1500 x g. Subsequently, the red and white blood cells are stacked at the bottom of the tube under the separator gel, whereas the platelets were sedimented above the gel layer with the plasma floating above them. After a resuspension step, around 5.5 mL PRP per tube were obtained and stored until use for a maximum of 15 days at room temperature in a polypropylene tube (Becton-Dickinson). To obtain platelet-poor plasma (PPP), we collected the plasma directly after centrifugation without the resuspension step.

Platelets, red and white blood cells, as well as mean platelet volume were counted (Sysmex) in whole blood before centrifugation and in the prepared PRP/PPP before addition to culture media. In all PRP/PPP conditions, heparin (Leo) was added (2UI/ml) to prevent clot formation.

### Cell proliferation assays

GF proliferation was measured by using a ready-to-use solution containing stable tetrazolium salt WST-1 (Cell Proliferation Reagent WST-1, Sigma-Aldrich). Ninety-six well plates were seeded with 5 x 10^3^cells and incubated for 2 hours in GM at 37°C in 5% CO2 humidified atmosphere. GM with nicotine ((-)-Nicotine, Sigma-Aldrich) at different final concentrations (25 ng/ml, 100 ng/ml, 250 ng/ml, 500 ng/ml) was added to each well. The medium was replaced after 48 h and complemented with 10% PRP. After 2 and 5 days of incubation, 10 µl of Cell Proliferation Reagent WST-1 were added to each well and the plate was incubated for 4 h before the measure of the absorbance of the formazan at 460 nm in a Cytation plate reader (Biotek).

### Crystal violet viability assay

GF were plated at a density of 5 x10^3^cells per well in 96-well plates. Cells were treated with various final concentrations of nicotine (25, 100, 250 and 500 ng/ml) in GM. After 48 h, the medium was replaced and complemented with 10% PRP. After 2 and 5 days of incubation, cells were fixed with 100 µl of methanol at -20°C for 15 min. A solution of 0.1% crystal violet (100 µl) was added to each well for 10-15 min, washed with distilled water and air-dried. Cytation reader was used to image the plates. Samples were then dissolved with glacial acetic acid, and the optical density at 570 nm was determined using an automatic microplate reader (Biotek).

### CellTiter-Glo ® luminescent Cell cell viability assay

GF were plated in GM at a density of 5 x10^3^ cells per well in 96-well plates for 2 hours to allow cell attachment. Culture media with nicotine [25-500 ng/ml] was used for 6 days, and 10 % PRP- was added to the nicotine culture media for 3 more days. Cell viability was assessed in accordance with the manufacturer’s instructions. Luminescence was recorded on a Cytation multi-mode reader in relative light units (RLU).

### Live/Dead test

GF were seeded at a density of 5 x 10^3^ cells in 96-well plates and incubated for 2 h at 37°C in 5% CO_2_ humidified atmosphere. DMEM with nicotine ((-)-Nicotine, Sigma-Aldrich) at different final concentrations (25 ng/ml, 100 ng/ml, 250 ng/ml, 500 ng/ml) was added to each well. After 48 h, 10% PRP were added to the different conditions. At the end of 2 days of incubation, Live/Dead assay (Invitrogen) was performed following standard protocol of manufacturer to determine viability of GF cells. The discrimination of live cells from dead cells was realized by respectively and simultaneously staining with green fluorescent calcein to indicate esterase activity and red-fluorescent ethidium bromide to indicate loss of plasma membrane integrity by visualization on Cytation. Gen 5 software was used to quantify the amount of calcein positive cells in the wells.

### Wound migration assay

GF at a density of 4 × 10^5^/ml (70 μl volume) were added to 24-well plates with culture insert (Ibidi). The cells were incubated at 37 °C for 24 h to allow cell attachment. Then, a cell-free gap of 500 μm was created by removal of the culture insert. For the measurement of cell migration, 250 μL of mitomycin C (Thermo Fisher Scientifics) at 0.4 mg/ml was added to each well to avoid cell proliferation. Medium was replaced by test media in triplicates: DMEM, heparin (2 IU/ml) supplemented with 10 % FBS, 10 % PRP, nicotine 250 ng/ml, nicotine 500 ng/ml alone or combined with 10% PRP. Images were captured after 24 h using an inverted phase-contrast microscope at 4× magnification. The percentage of wound closure was calculated with ImageJ software.

### Chronicity and senescence test

GF cells were seeded at low densities (2 x 10^3^) in 35 mm Petri dishes (Falcon) and incubated for 2 h at 37°C in a 5 % CO2 humidified atmosphere. DMEM with nicotine ((-)-Nicotine, Sigma-Aldrich) at different final concentrations (25, 100, 250 and 500 ng/ml) was added to each well. When the cells were confluent, they were passed to other 35 mm Petri dishes using TrypLE (Gibco). This procedure was repeated weekly over a period of one month until P10. Cell seeding concentrations decreased over time due to cell damage with 250 and 500 ng/ml nicotine experimental conditions. GF cells were fixed and stained following the protocol of senescence associated (SA) β-Galactosidase Staining kit (Cell Signaling Technology), designed to detect β-galactosidase activity, a known characteristic of senescent cells. Quantification was made with ImageJ software by measuring the total cell surface and counting the SA-β-galactosidase positive cells.

### DCFDA cellular ROS assay

Intracellular reactive oxygen species (ROS) was assessed using the DCFDA Cellular ROS Detection Assay Kit (Abcam), according to the manufacturer’s protocol and as previously described [16]. GF were cultured in 96 well plates (1 x 10^4^ cells/well) for 48 h with increasing concentrations of nicotine (50-500 ng/ml) and then 10% PRP was added for 3 days. The concentrations of the positive control [tert-butyl hydroperoxide (TBHP)] was 200 μmol/l. Fluorescence at 483-14 nm/530-30 nm was recorded from 3 h up to 24 h post-treatment, using the Cytation microplate reader. Background fluorescence was subtracted from each value before fold increase in fluorescence intensity relative to the negative control (GM) was determined.

### Acridine orange staining

Cells were seeded in 24-well microclear optical plates (1 x 10^4^ cells/well). After 24h, they were incubated with increasing concentrations of nicotine for 48 h, followed by 10% PRP addition for 3 more days. Then, the cells were stained with medium containing 1 μg/ml acridine orange for 15 min at 37 °C, washed twice in PBS and immediately observed with a Cytation imager. The cytoplasm and nucleus of acridine orange-stained cells fluoresced bright green, whereas the acidic autophagic vacuoles fluoresced bright red, as described previously [17, 18].

### *C. elegans* strains and maintenance

Standard methods for culturing and handling *C. elegans* were followed [14]. *C. elegans* was maintained on standard nematode growth medium (NGM) plates streaked with OP50 *E. coli*. The strains used in this study were obtained from the *C. elegans* Genetics Center (University of Minnesota, MN, USA). Animals were transferred to fresh NGM plates without floxuridine (FUdR) every 2 days to ensure maintenance. The strains used in this study include the N2 Bristol strain and MAH215 sqIs11[lgg-1p::mCherry::GFP::lgg-1 + rol-6].

### Lifespan in N2 wild-type *C. elegans* with PRP and PPP

Animals at larval stage 4 (L4) were divided into triplicates in a 96-well plate, with 10 worms per well in 150 µL of liquid culture medium (S Basal, 10 mM potassium citrate pH 6, 10 mM metal solution, 30 mM CaCl₂, 30 mM MgSO₄, 50 µM FuDR) supplemented with OP50 [19] Freshly prepared platelet-poor plasma (PPP) or PRP was added at 10% of the total volume in designated wells, along with heparin (2UI/ml) [20].The animals were counted every other day, and the medium was refreshed every 2 days. Graphs were generated using GraphPad Prism 9.

### Confocal microscopy in MAH215 sqIs11 *C. elegans*

L4 *C. elegans* were synchronized [14] and deposited (10 worms) in 150 µL of liquid culture medium (S Basal, 10 mM potassium citrate pH 6, 10 mM metal solution, 30 mM CaCl₂, 30 mM MgSO₄, 50 µM FuDR) with OP50 *E. coli* bacteria in a 96-well plate [19]. Nicotine titration was performed over 4 days by adding 10 ng/mL, 100 ng/mL, or 1 µg/mL of nicotine to the liquid culture. To assess the effect of PRP on autophagy, 10% freshly prepared PPP or PRP was added to the culture medium, along with heparin (2UI/ml). Before imaging, worms were placed on a drop of 60% glycerol on a 2% agarose pad and imaged using a Zeiss LSM800 confocal microscope. Autolysosomes quantifications were performed in the anterior part of the intestine, where autophagy is most prominent, using QuPath software. The mean fluorescence intensity of the selected area is represented.

### Angiogenesis Protein Array in GF

RayBio Human Angiogenesis Antibody Arrays C-1000 (RayBiotech) were used to assay the GF-secreted cytokines in conditioned medium containing 10% PRP, Nicotine (25-100-250-500 ng/ml) or a combination of both. To prepare the conditioned medium, GF were seeded in T25 flasks at a concentration of 5 × 10^5^ cells per flask in complete growth medium. The following day, the medium was replaced with DMEM and 0.5% FBS, and the cells were treated with the different experimental conditions After 48 h, the medium was collected and centrifuged at 400× g for 5 min to remove cellular debris. The membranes were incubated with fresh supernatants. In the array, 30 angiogenic cytokines, chemokines, enzymes and growth factors were measured according to the manufacturer’s protocol. Membranes were imaged using a Fusion System (Vilber Lourmat). Protein levels were quantified using the Protein Array Analyzer in the ImageJ program. Values were normalized to reference spots on the membranes.

### Statistics

Data are presented as mean ± SD, and statistical differences were determined using the statistical test specified in the figure legends, with * p < 0.05, ** p < 0.01, ***p < 0.001, and **** p < 0.0001. For the comparison of two populations, a t-test or Mann–Whitney test was used depending on the normality of the dataset. For the comparison of more than two populations, either ANOVA or the Kruskal–Wallis test was used, depending on the normality of the dataset.

## Results

### Nicotine induces morphological changes in human gingival fibroblasts that are counterbalanced by PRP treatment

At 48 h exposure, low amount of nicotine (25, 100 ng/ml) did not induce visible morphological changes in GF, whereas the two highest doses of nicotine (250 and 500 ng/ml) brought an increase in vacuolization. The addition of 10% PRP for another 48 h reversed the phenotype and led to an increase in cell number (Figure 1).

**Figure 1.**
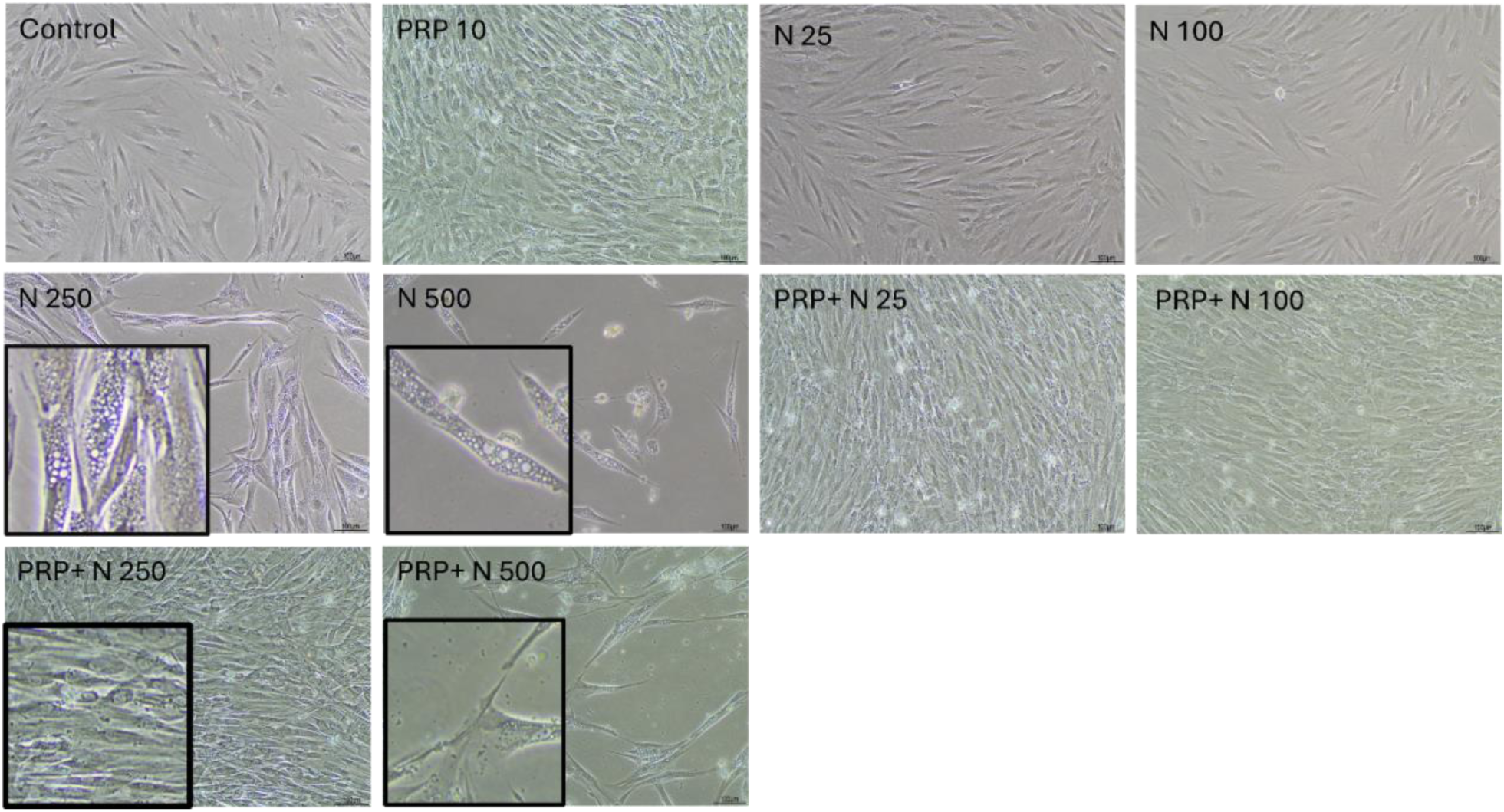
Nicotine-induced phenotypic modifications in GF are reversed by PRP treatment. Increasing concentrations of nicotine (25, 100, 250, 500 ng/ml) were added to GF culture for 4 days. At Day 2, PRP was added in different conditions. Bright-field pictures at 10x. Boxes are zoomed-in magnifications.

### High doses nicotine exposure led to significant alterations in the metabolic activity and viability of GF, which were partially reversed by PRP

Nicotine induced a significant decrease in metabolic activity after 96 h treatment with 500 ng/ml as evaluated by WST-1 incorporation (Figure 2A). In control condition with heparin, the metabolic activity was not significatively different than in control condition without heparin showing that 2UI/ml heparin had no effect on the GF. Hence, heparin alone controls were not shown in the following experiments. Heparin-added 10 % PRP without nicotine induced a potent effect on cell proliferation, metabolism and viability (Figure 2A, 2B, 2D, 2F).

**Figure 2.**
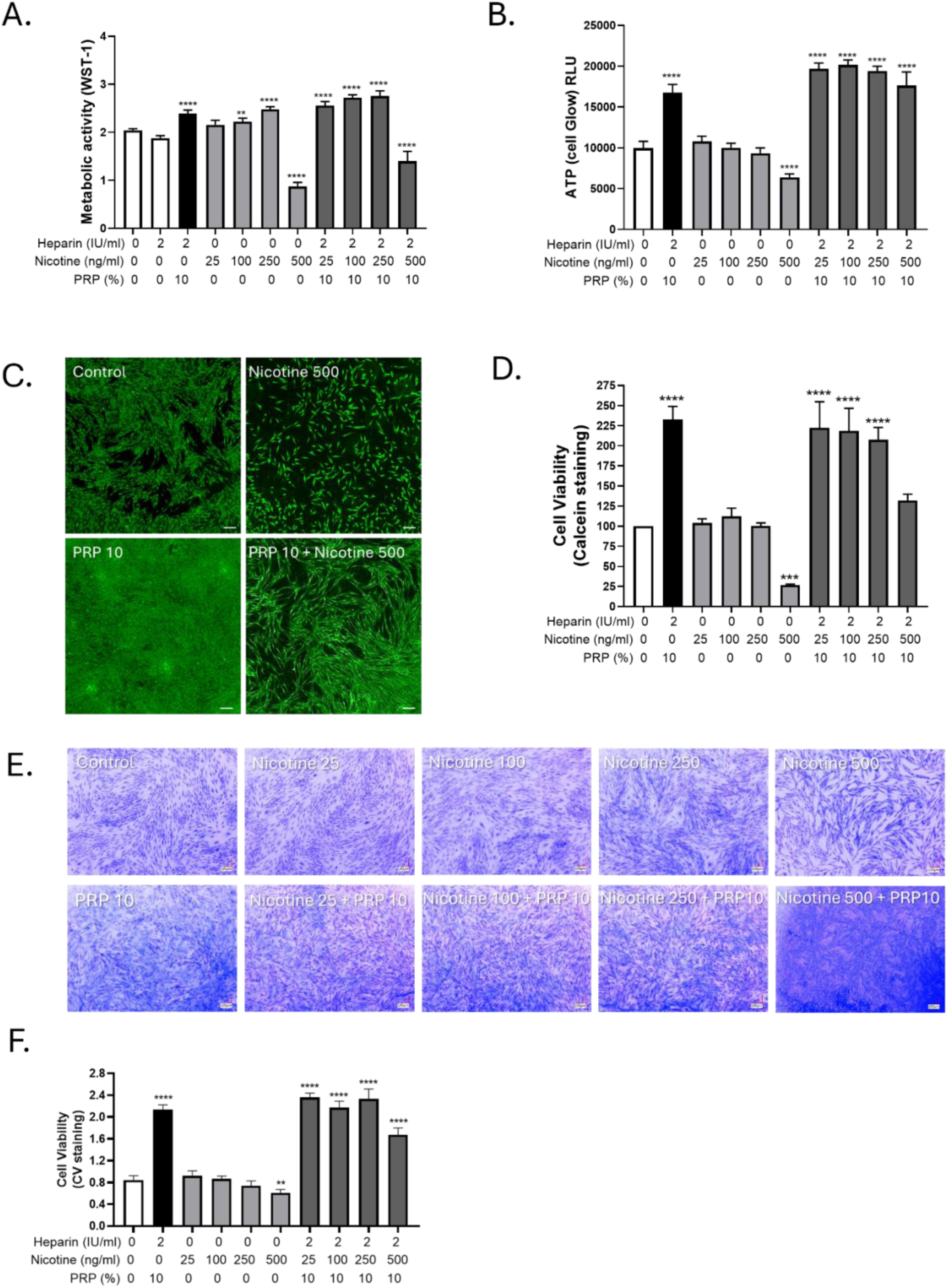
Nicotine impaired GF metabolic activity and viability, mitigated by PRP. Increasing concentrations of nicotine (25, 100, 250, 500 ng/ml) were added to GF culture for 4 days. At Day 2, PRP was added in the annotated conditions. Heparin alone was used as the control for PRP conditions. A. Metabolic activity of the cells assessed by WST-1 incorporation. B. Total intracellular ATP production assessed by Cell Titer Glow. C. Calcein microscopic acquisitions at 4x (scale bar= 100 µm). D. Cell viability is assessed by calcein incorporation followed by fluorescent measurement. E. Crystal violet for cellular viability assessment, pictures taken at 10x magnification. F. Quantification of crystal violet dye release by absorbance measurement. Data presented in this figure are from representative experiments from at least 3 independent repetitions. **p < 0.01, ***p < 0.001 and ***p <0,0001 compared to control condition.

Consistently, the intracellular ATP production measured via Cell Titer Glow assay (Figure 2B) was also significantly decreased in cells treated with the highest concentration of nicotine, indicating impaired cellular energy metabolism. Fluorescent microscopy of calcein-stained cells (Figure 2C) further supported these findings, with visibly reduced green fluorescence in high nicotine-exposed conditions, reflective of compromised cell viability. Quantitative calcein incorporation (Figure 2D) confirmed a significant decrease in viable cells compared to untreated controls (p < 0.05). In parallel, crystal violet staining (Figure 2E) demonstrated reduced cell density in nicotine-treated cultures, with absorbance quantification (Figure 2F) corroborating a significant loss in cell number. Notably, the addition of 10% PRP after two days of nicotine exposure partially rescued both metabolic and viability metrics across all assays. These data indicate that high dose nicotine negatively impacted GF metabolic function and survival, while PRP exerted a protective effect, enhancing recovery from nicotine-induced cytotoxic stress.

### PRP restored cell migration impaired by nicotine exposure in GF

Representative images from the Ibidi chamber assay (Figure 3A) with visibly wider gaps remaining in control groups and more complete closure in nicotine alone, PRP alone and nicotine-PRP-rescued conditions. Together, these results indicate that 10% PRP exerted a synergistic effect together with nicotine (250 ng/ml) on fibroblast migration, highlighting its potential role in supporting wound healing in nicotine-compromised environments. PRP and nicotine together significantly improved wound closure, suggesting a pro-migratory effect.

**Figure 3.**
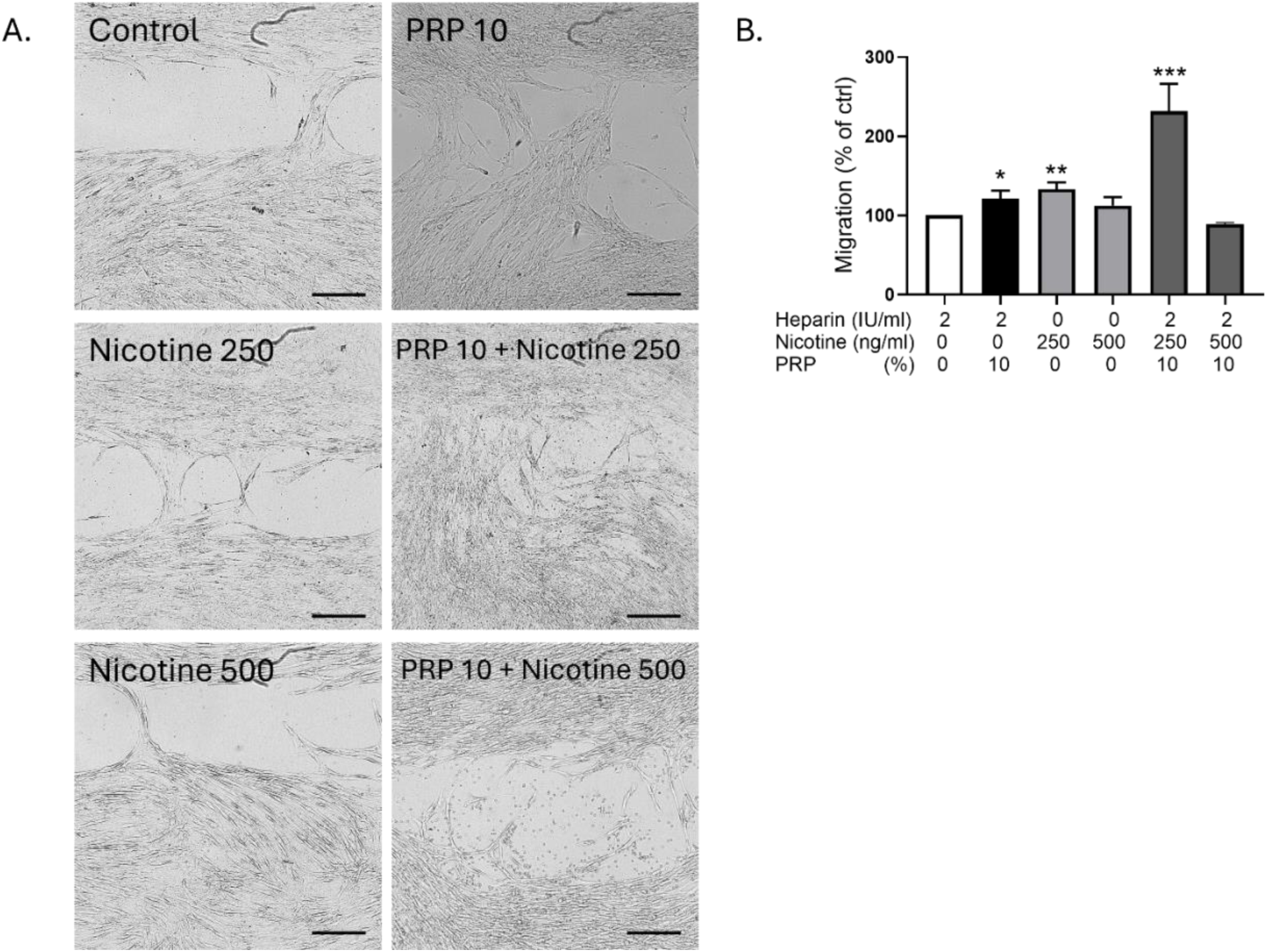
Effect of nicotine and PRP on the migration of GF. A. Wound healing pictures of Ibidi chambers after 24 h of GF migration in the presence of the various treatments. B. Wound Healing quantification: areas measured at Day 0 divided by area measured after 24 h of migration. N=4 pictures per experimental condition. *p<0.5, **p < 0.01 and ***p < 0.001.

Wound healing quantification (Figure 3B) demonstrated a slight but significant increase in gap closure in nicotine-treated cells (250 ng/ml) and in 10% PRP treated cells after 24 h compared to control conditions, indicating that nicotine enhanced GF migration at that concentration.

### PRP reduced nicotine-induced cellular senescence in GF

*In vitro* senescence was induced by serial passaging of GF at low density from passages 2 to 7, followed by treatment with nicotine and/or PRP. At passage 8, senescence-associated β-galactosidase (SA-β-gal) staining revealed a marked increase in β-gal–positive cells in the nicotine-treated group compared to control, as visualized microscopically (Figure 4A). Quantitative analysis (Figure 4B) showed a significant elevation in the proportion of β-gal–positive cells under nicotine exposure, indicating enhanced senescence. In contrast, PRP treatment significantly reduced the percentage of β-gal-positive cells, both in nicotine-treated and control conditions, demonstrating its anti-senescent effect. The most striking effect was that the 500 ng/ml nicotine induced total cell death that was prevented by 10% PRP addition showing B-gal values even lower than in control condition. These results suggest that PRP mitigates nicotine-induced senescence in GF, supporting its potential regenerative and protective role in maintaining fibroblast function.

**Figure 4:**
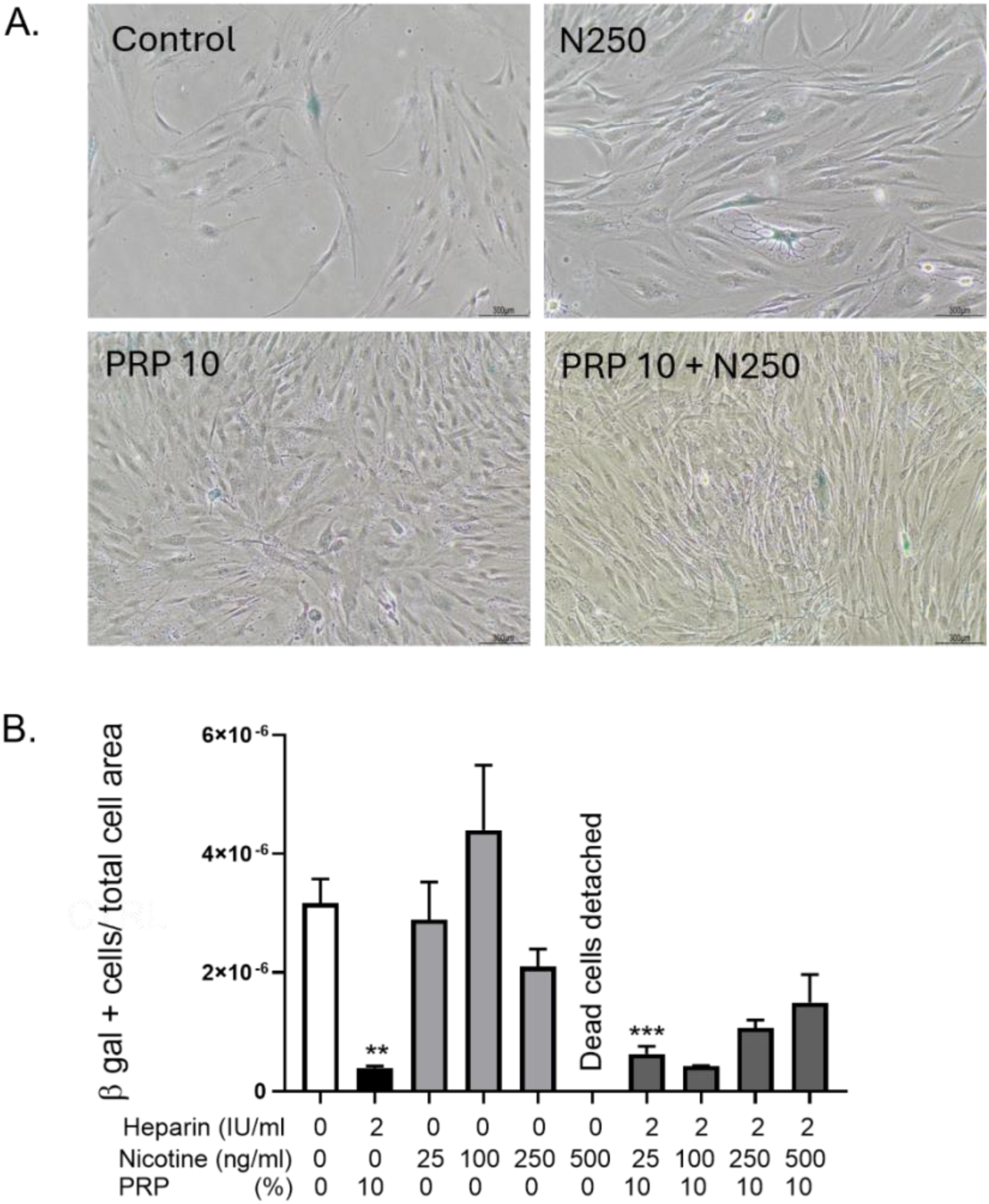
PRP exerted an anti-senescence activity on GF. To evaluate the effect of nicotine and PRP on GF senescence, we induced *in vitro* aging by cultivating the cells at a low density from passages 2 to 6. A. Senescence associated β-galactosidase (SA-beta gal) staining was performed on GF at passage 8 (10x objective lens). B. Quantification of β-gal positive cells reported to the total cell number. *p<0.5, **p < 0.01 and ***p<0.001.

### Nicotine treatment induced a significant increase in intracellular ROS levels in GF that was neutralized by PRP

Fluorescence microscopy revealed enhanced green fluorescence intensity in nicotine-treated cells compared to control, indicating elevated ROS production (Figure 5A). Quantitative analysis of fluorescence per field (Figure 5B) showed a statistically significant increase in ROS levels following nicotine exposure (*p* < 0.01), while co-treatment with PRP markedly reduced ROS accumulation. These findings demonstrate that nicotine promoted oxidative stress in fibroblasts, and that PRP exerted a protective antioxidant effect, attenuating ROS generation.

**Figure 5:**
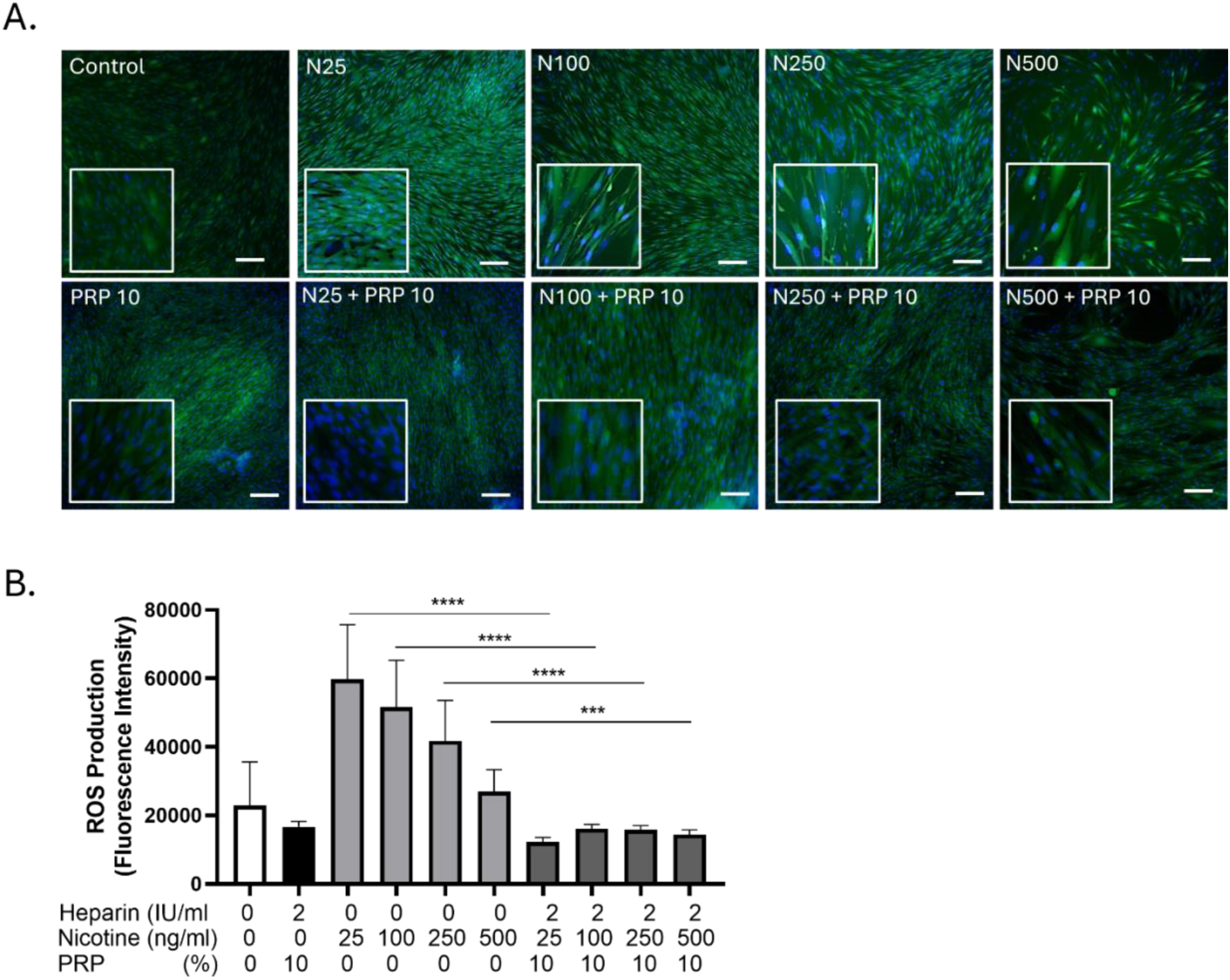
PRP prevented intracellular ROS induced by nicotine in GF. A. Fluorescence imaging: Representative confocal/epifluorescence micrographs of GF after DCF-DA staining (green). Nuclear counterstaining with Hoechst (blue); scale bar = 10 µm. B. Quantification: Average fluorescence intensity per field. Data are expressed as mean ± SEM. Unpaired t test, ***p < 0.001, ****p<0.0001.

### Nicotine exposure increased the formation of acidic vesicular organelles in GF, an effect that was attenuated by PRP

After 24 h exposure of GF to nicotine at 250 or 500 ng/mL, acridine orange staining revealed a marked increase in yellow fluorescence, indicating an accumulation of acidic compartments such as lysosomes or autophagic vacuoles (Figure 6A). An acridine orange signal was spotted at 250 and 500 ng/mL nicotine. In contrast, co-treatment with 10% PRP markedly reduced the fluorescence, suggesting a restoration of lysosomal homeostasis and/or reduced autophagic stress. Hoechst nuclear staining (blue) confirmed proper subcellular localization and overall cell integrity across conditions. These results indicate that nicotine disrupted vesicular acidification in GF, while PRP counteracted these effects, supporting its role in maintaining cellular homeostasis under oxidative or stress-inducing conditions as shown by the quantification (Figure 6B).

**Figure 6:**
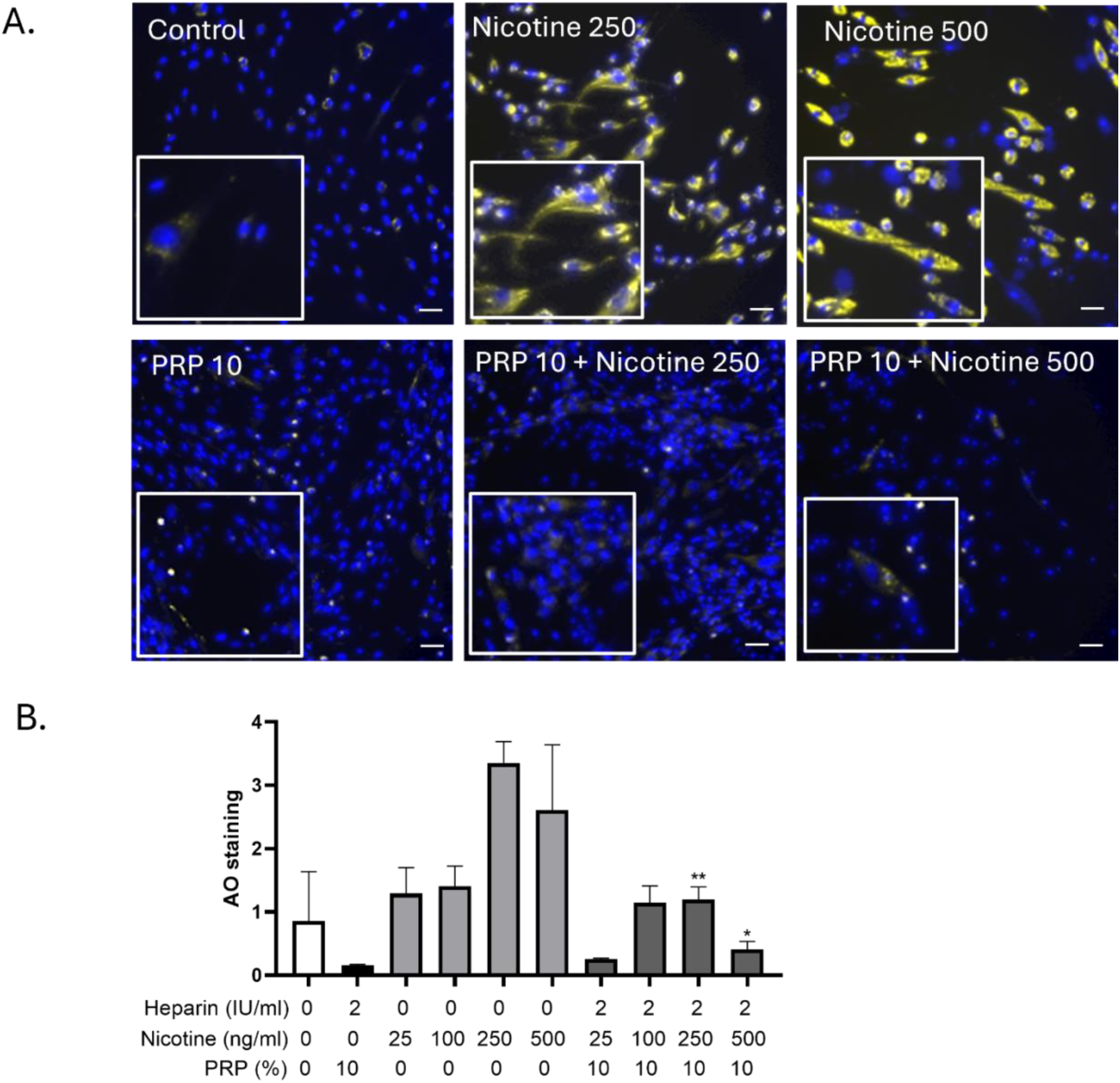
Autophagy evaluation by acridine orange staining of GF following nicotine and/or PRP exposure. GF were treated with nicotine (250 and 500 ng/ml) and PRP 10% as indicated for 24 h. Acridine orange staining (1 µM, 20 min) was evaluated by fluorescence microscopy, as well as Hoechst nuclear dye (blue) to depict subcellular localization.

### PRP conferred significant protective effects against nicotine-induced premature aging in C. elegans

Lifespan analysis of wild-type N2 worms in liquid culture revealed a reduction in survival along time due to aging, which was partially mitigated by treatment with Plasma-Poor-platelets (blue line, *p* < 0.05), and more robustly by PRP (red line, *p* < 0.01), indicating a dose-dependent protective effect of blood-derived plasma components (Figure 7A). To further assess the impact on cellular aging processes, autophagic flux was evaluated using the mCherry::GFP::LGG-1 reporter strain. Quantification of autolysosomes *via* mCherry fluorescence showed a significant increase in autophagic activity in worms exposed to increasing concentrations of nicotine for 4 days (Figure 7B), reflecting stress-induced autophagy. However, co-treatment with 10% PRP during the final 3 days significantly reduced autolysosome accumulation (Figure 7C, white bar vs. grey bar), suggesting PRP decreased excessive autophagy linked to nicotine-induced stress. Representative fluorescence images support these findings, displaying visibly reduced mCherry puncta in the PRP-treated group. Collectively, these data indicate that PRP mitigated nicotine-induced lifespan shortening and aberrant autophagic responses in *C. elegans*, underscoring its potential anti-aging and cytoprotective properties.

**Figure 7:**
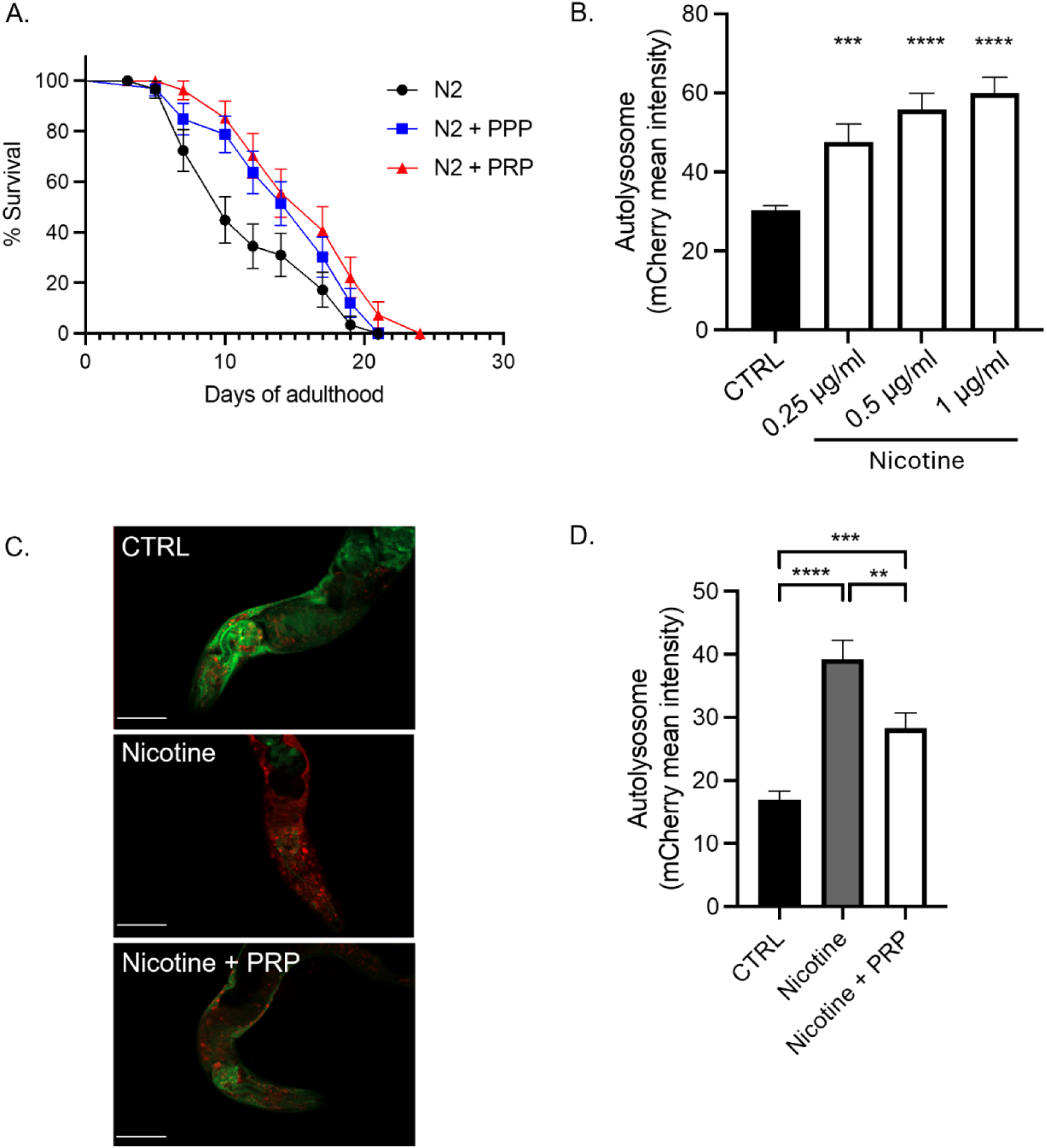
PRP conferred protective effects against nicotine-induced premature aging in *C. elegans*. A. Lifespan assay on N2 wild type *C. elegans* in liquid culture. PPP (blue line, p < 0.05) or PRP (red line, p< 0.01) treatment at 10% were compared to the control (black line). B. and C. Autolysosome quantification was realized by measuring mCherry fluorescence from the mCherrry::GFP::LGG-1 strain treated with (B) different concentration of nicotine for 4 days or (C) with 1 μg/ml of nicotine for 4 days (grey bar), in addition to PRP at 10% for the last 3 days (white bar). One representative image of a *C. elegans* in each condition is depicted, where the scale bar represents 50 μm (C). D. Standard error of mean is represented for each graph. The Log-rank Mantel-cox test was realized for the statistics of the lifespan, and an unpaired t-test was realized for the autolysosome quantification.

### Profiling of the cytokine secretome by the GF in the presence of increasing concentrations of nicotine, PRP or a combination of both

To further identify the relevant mediator(s) responsible for cellular deleterious effects of nicotine and the regenerative activity of PRP, a cytokine antibody array was performed.

Growth factors released from GF were analyzed, while cultivated with nicotine (25-500 ng/ml), PRP (10%) or a combination of both. The array was performed on the proteins released from GF subjected 3 days to nicotine and then treated for 3 more days in PRP (Figure 8A). Analysis (heat map arrays) of proteins secreted are shown in Figure 8B. A uniform pattern of modulation among the cytokines spotted was observed. An increase of ANG, GRO, Il-6, IL-8, MCP-1, RANTES, TIMP-1 and TIMP-2 in the conditions where PRP is present was depicted, with the most predominant secretion at N25 + PRP 10 for all cytokines. The presence of VEGF-A in the secretome was also observed, however its secretion and modulation were lower than the other cytokines. Examples of four angiogenesis cytokine antibody array membranes for GF cultures with heparin, PRP 10%, Nicotine 500 ng/ml or PRP 10 % and Nicotine 500ng/ml that were used for quantification (Figure 8C).

**Figure 8:**
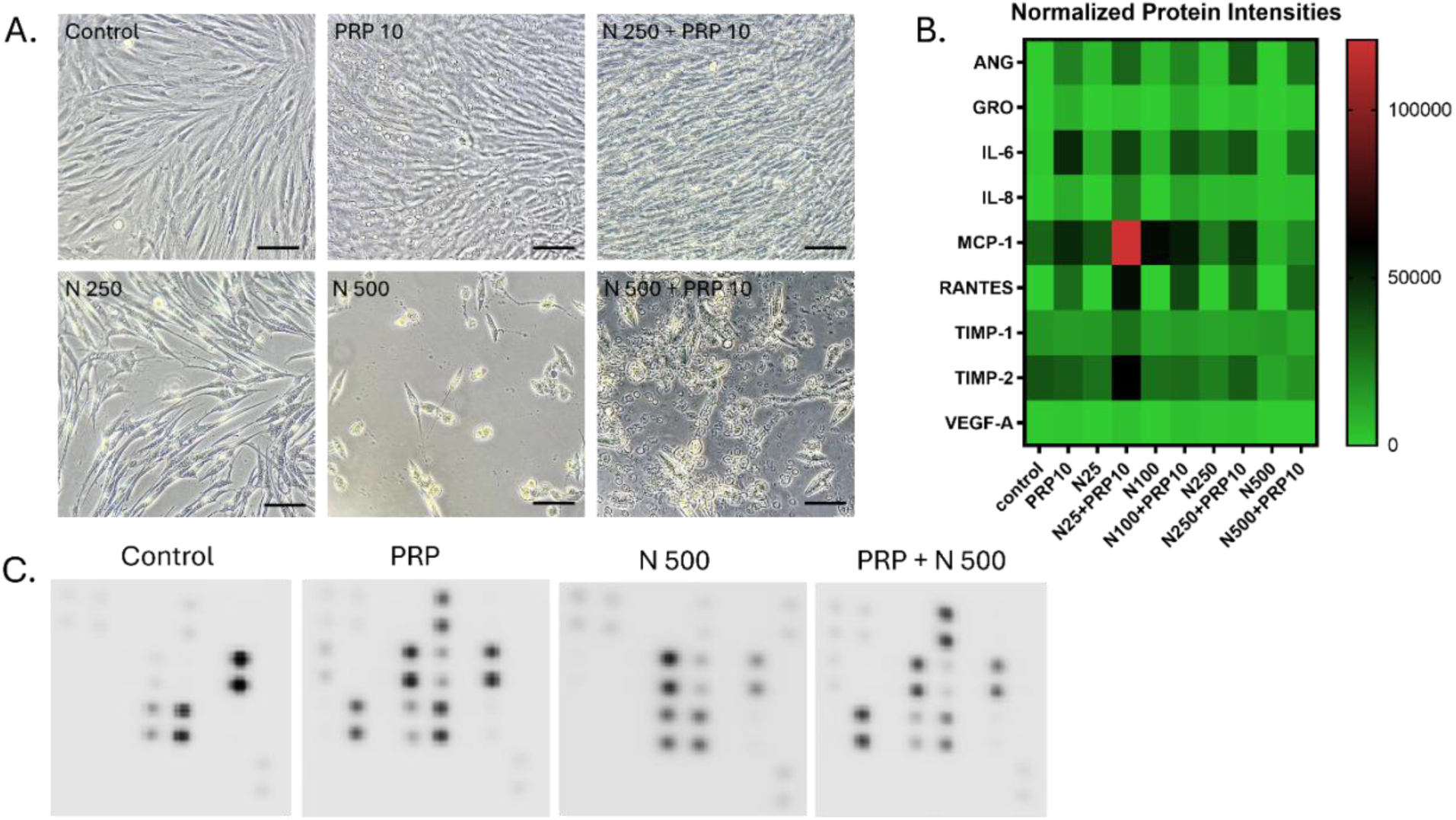
Secretome profiling of cytokines present in the conditioned medium of GF in Nicotine, PRP or both combined. A. Representative images of the cultures used to produce conditioned medium for secretome profiling on cytokine arrays. Scale bars: 300 μm. B. Heat map analysis representing the average pixel density of the duplicated spots for each cytokine highlighted the array quantified by densitometry. Variation in color intensities represents differential secretion of cytokines by GF, in the presence of PRP, Nicotine (25-500 ng/ml), or the combinations of both. C. Example of the human angiogenesis antibody array composed of duplicated spots of 23 angiogenic and inflammation-related factors performed with the culture supernatants.

## Discussion

In the present study we aimed to evaluate the effect of PRP on several biological activities of nicotine-treated GF and *in vivo* models of *C. elegans*. We made the hypothesis that PRP could compensate for the negative effects of nicotine on the cells of the periodontium and could therefore be beneficial in the treatment of periodontitis in smokers. Our results showed that PRP has the capacity to modulate several key cellular processes affected by smoking.

Smoking is a major etiological factor in the development and progression of periodontal diseases and is known to impair the outcomes of periodontal therapy [21, 22]. One of the primary cellular targets affected by tobacco smoke are GF, which play a critical role in extracellular matrix production, wound healing, and immune modulation [21]. Nicotine, a key alkaloid in tobacco, has been shown to disrupt fibroblast functions through multiple cytotoxic mechanisms, including oxidative stress, autophagic dysregulation, and pro-inflammatory signaling [23, 24].

Our findings confirm and extend prior studies demonstrating that nicotine induced a range of detrimental effects in human GF. These include decreased metabolic activity and viability, impaired cell migration, increased production of reactive oxygen species (ROS), elevated lysosomal activity, morphological changes, and premature cellular senescence, each of which is known to contribute to impaired periodontal healing [25].

Previous literature demonstrated that nicotine exhibited biphasic effects, with low concentrations promoting proliferation in mesenchymal stem cells [26], while higher concentrations tend to be cytotoxic, inhibiting proliferation, protein synthesis, and attachment [27]. In our study, clinically relevant nicotine concentrations (50–500 ng/mL; ≈0.3–3.0 μM) were chosen to reflect systemic exposure levels in smokers [28, 29]. These concentrations were sufficient to elicit significant cellular dysfunction, highlighting that even low-level chronic exposure could impair periodontal health.

High nicotine concentrations were previously shown to inhibit collagen and fibronectin synthesis [30], reduce alkaline phosphatase activity [30], and stimulate ROS generation leading to mitochondrial damage [31]. A significant increase in intracellular ROS in nicotine-treated GF was confirmed in the present study, supporting nicotine’s role as a potent oxidative stressor. Similarly, the increase in acidic vesicular organelles, indicating autophagic activity, consistent with previous observations in cardiomyocytes [32]. Previous reports showed that nicotine could destabilize lysosomes, possibly through oxidative mechanisms [33]. Furthermore, our study confirmed nicotine-induced senescence in GF via β-gal staining, in line with reports of accelerated aging in lung and skin fibroblasts exposed to tobacco constituents [34]. This premature senescence may contribute to the observed impairment in fibroblast proliferation and tissue regeneration.

Platelet-Rich Plasma (PRP), a preparation containing concentrated platelets suspended in plasma, has shown increasing promise in tissue engineering and regenerative medicine due to its rich content of growth factors such as platelet-derived growth factor (PDGF), vascular endothelial growth factor (VEGF), and transforming growth factor (TGF)-β [35, 36]. Our findings showed that 10% PRP supplementation in culture media significantly increased cell proliferation, but also restored metabolic activity, improved migration, and mitigated ROS generation, lysosomal stress, and cellular senescence in nicotine-exposed GF. PRP may exert these cytoprotective effects *via* multiple pathways. TGF-β and PDGF are known to modulate the mechanistic Target of Rapamycin (mTOR) and SMAD signaling cascades, both of which intersect with autophagy regulation and also have homologs in *C. elegans* [37, 38].

Our results show that by restoring autophagic flux, PRP helped maintain cellular homeostasis and prevented nicotine-induced toxicity. Other studies have shown that PRP reduced Interleukin (IL)-6 and Tumor Necrosis Factor (TNF)-α levels, likely through inhibition of Nuclear Factor (NF)-κB signaling—thereby attenuating inflammation [39].

The autophagy process is crucial for survival across species, and it has been shown that autophagy becomes less effective with aging, especially in *C. elegans*. For this reason, our study was conducted on age-synchronized young nematodes. The dual modulation of autophagy and inflammation by PRP aligned with reports that PRP-treated fibroblasts exhibited enhanced collagen synthesis, migration, and extracellular matrix remodeling [40]. Our data further suggest that PRP displayed an anti-senescent activity, a previously underexplored mechanism in periodontal applications. In *C. elegans*, nicotine triggered a hyperactivation of autophagic processes, leading to a state resembling to an aging worm [41]. PRP exposure reduced autolysosomal accumulation and extended lifespan under nicotine stress, underscoring the evolutionary conservation of its cytoprotective effects.

Cytokine protein profiling in the secretome of GF in the presence of nicotine and PRP revealed that cytokines MCP-1, IL-6, RANTES/CCL-5 and TIMP-2 are increased. This is particularly interesting because these four cytokines are linked to autophagy. TIMP-2 is an inhibitor of matrix metalloproteases (MMPs) and regulates extracellular matrix degradation and angiogenesis. Inpancreatic stellate cells, it has been shown that when autophagy is inhibited, TIMP-2 levels decrease while MMPs increase [42]. MCP-1 is a key chemokine involved in recruiting immune cells to inflammatory sites. Its role in autophagy has been described [43, 44] and notably p62, a key autophagy-related protein, regulates MCP-1 [45]. A link between increased autophagy and elevated cytokines, including MCP-1 and IL-6, has also been reported in synovial fluid neutrophils [46]. Interleukin-6 (IL-6) is a pro-inflammatory cytokine. It activates the JAK/STAT3 pathway and consequently inhibits autophagy [47]. RANTES/CCL5 recruits immune cells, activates mTOR, and thereby inhibits autophagy [48, 49]. It is plausible that RANTES/CCL5, like IL-6, is overexpressed after nicotine + PRP treatment to dampen autophagy and avoid runaway activation. Consequently, nicotine exposure in GF induces autophagy, which PRP tends to reduce to prevent autophagy-dependent cell death, also known as autosis.

These findings have significant relevance for periodontal therapy in smokers. Nicotine-induced impairment in GF function contributes to delayed healing, reduced tissue regeneration, and greater susceptibility to chronic periodontitis [24, 50, 51]. PRP’s ability to counteract these effects makes it a compelling adjunct in regenerative dentistry, particularly for patients with tobacco exposure. Additionally, dissecting the contributions of individual PRP components (e.g., IGF-1, VEGF, antioxidant enzymes) through proteomic and transcriptomic analyses would further clarify the mechanisms underlying its therapeutic efficacy.

From a clinical point of view, a recent systematic review [52] underscored the potential of PRP as a valuable adjunct in oral surgery, demonstrating significant benefits in the regeneration of soft tissues and, to a lesser extent, hard tissues with no distinction between smokers and non-smokers.

However, translation to clinical practice warrants caution. Our study was limited to *in vitro* studies using primary fibroblasts derived from non-smokers, as well as invertebrate models. Further investigations should employ *ex vivo* tissue explants from smokers and non-smokers (e.g., 3D Oral Mucosa Constructs) that are closer to clinical reality than 2D cell culture.

Another interesting point would be to treat those constructs with autologous PRP to evaluate eventual systemic alterations in PRP from smokers that would lower its regenerative potential.

## Acknowledgements

The *C. elegans* strains were kindly provided by the *C. elegans* Genetics Center (University of Minnesota, MN, USA). A great thanks to Nicolas Liaudet, who coded the script for picture quantification on QuPath and to the Bioimaging facility of the UNIGE for their help.

